# Yield losses associated with peanut smut incidence in Argentina: a quantitative synthesis across field studies

**DOI:** 10.64898/2026.08.11.744131

**Authors:** Luis I. Cazón, Noelia R. González, Emerson M. Del Ponte, Ana C. C. de Carvalho, Florencia Asinari, Boris X. Camiletti, Juan A. Paredes

## Abstract

Peanut smut, caused by *Thecaphora frezzii*, is an important constraint to peanut production in Argentina, but quantitative estimates of yield losses across environments remain limited. We quantified the relationship between disease incidence and kernel yield using 922 observations from 26 field studies conducted in Córdoba, Argentina, between 2021 and 2025. Study-specific incidence–yield relationships were analyzed using linear regression, random-effects meta-analysis, and linear mixed-effects models. Peanut smut incidence was consistently associated with yield reduction across studies. The estimated damage coefficient ranged from 24.2 to 28.7 kg ha⁻¹ per 1% increase in disease incidence, corresponding to a relative yield reduction of 0.74–0.87% of attainable yield. In contrast, attainable yield varied markedly among studies, ranging from 1,370 to 5,409 kg ha⁻¹. Although an exploratory segmented analysis suggested a breakpoint near 12% incidence, subsequent moderator analyses, study-specific regressions, and normalized response curves provided no evidence of a biologically meaningful change in the damage coefficient across incidence or yield classes. These results indicate that differences among environments were primarily associated with attainable yield rather than with changes in the magnitude of disease-associated yield loss. The resulting damage function provides a quantitative basis for yield-loss assessment and disease management in peanut.

## INTRODUCTION

Peanut smut, caused by the soilborne pathogen fungus *Thecaphora frezzii* Carranza & Lindquist, is currently considered the most important disease affecting peanut (*Arachis hypogaea* L.) production in Argentina (Cazón and Paredes, 2026; Paredes et al., 2024; Rago et al., 2017). Since its first report in commercial fields in 1995 (Marinelli et al., 1995), the disease has rapidly established itself across the province of Córdoba, the main peanut-producing and exporting region of the country, reaching 100% field prevalence after approximately 15 years (Bonessi et al., 2011; Cazón et al., 2018; Cazzola et al., 2012; Rago et al., 2017). This rapid spread, combined with the pathogen’s persistence in soil and the limited effectiveness of current management strategies, has raised concern within the peanut production community.

*T. frezzii* survives in soil as teliospores, which can remain viable for extended periods in the absence of a host (Marinelli et al., 2008; Paredes et al., 2022). Infection occurs after flowering, when the fertilized gynophore penetrates the soil and releases exudates that stimulate teliospore germination, initiating the infection process (Astiz Gassó, 2008; Mary et al., 2020; van der Linde & Göhre, 2021). Fungal colonization leads to the formation of teliospores within developing pods, causing partial or complete destruction of seeds and potentially resulting in pod hypertrophy (Arias et al., 2021).

Because symptoms develop exclusively in underground pods, accurate assessment of peanut smut requires destructive sampling and visual inspection of mature pods (Cazón and Paredes, 2026; Paredes et al., 2022). Disease intensity is typically quantified using epidemiological parameters such as incidence and severity (Bock et al., 2021). For peanut smut, incidence is quantified as the proportion of infected pods, and severity describes the extent of damage within individual pods. In Argentina, pod severity is commonly assessed using a five-level diagrammatic scale, where level “0” represents completely healthy pods and level “4” corresponds to pods in which both kernels are entirely transformed into teliospore masses (Astiz Gassó et al., 2008). Given the near-proportional relationship between pod severity and incidence across disease intensities and susceptible cultivars, incidence was selected as the main response variable, as it consistently reflects the proportion of severely affected pods (Paredes et al., 2022; Paredes et al., 2024). This scale provides a standardized framework for disease assessment under field conditions and enables consistent comparisons across studies and production environments.

Peanut smut intensity varies across production regions, largely reflecting differences in cropping history and inoculum pressure. In the core production area of Córdoba, where peanut cultivation has been established for decades and processing facilities are concentrated, soils tend to be heavily infested with teliospores, resulting in higher disease levels. In contrast, recently expanded production areas in other provinces generally exhibit lower disease intensity (Paredes et al., 2022). This geographic variability is agronomically relevant because increased disease intensity has been associated with reduced crop productivity. Several studies have reported a negative association between peanut smut intensity and crop yield, with losses reaching up to 35% in highly affected fields (Paredes et al., 2020, 2022).

Despite this evidence, the quantitative relationship between disease incidence and yield loss has not been systematically evaluated across studies. Meta-analysis provides a robust framework for integrating results from independent experiments and identifying generalizable patterns (Del Ponte et al., 2024a). In other crop–pathogen systems, meta-analytic approaches have revealed robust disease–yield relationships despite substantial environmental variability, providing valuable information for yield-loss prediction, economic assessments, and disease management decision-making (Dalla Lana et al., 2015; Duffeck et al., 2020; Madden et al., 2009a; Yellareddygari et al., 2018).

Therefore, the objective of this study was to quantify the relationship between peanut smut incidence and crop yield using a meta-analytic mixed-effects modeling approach, aiming to improve the understanding of disease impact and to support the development of more effective management strategies.

## MATERIALS AND METHODS

### Data sources and database compilation

The database was assembled from previously published and unpublished field studies conducted by researchers at the Instituto Nacional de Tecnología Agropecuaria – Instituto de Patología Vegetal (INTA-IPAVE), Argentina. Field experiments were conducted between 2021 and 2025 in the core peanut-producing region of Córdoba Province, Argentina, in two production areas surrounding General Deheza and General Cabrera.

The compiled dataset comprised 922 observations derived from 26 independent studies, including 13 studies conducted in each production area. Each observation corresponded to a 1 m² field sampling area in which all harvested pods were evaluated for disease incidence and weighted to estimate the kernel yield (kg ha⁻¹). The experiments were originally designed to evaluate different management strategies for peanut smut control under field conditions and included measurements of disease intensity and crop yield, using commercially susceptible cultivars (ASEM 400, ASEM 450, and Granoleico). Disease incidence was expressed as the percentage of infected pods relative to the total number of pods evaluated (Rago et al. 2017).

To ensure consistency and statistical robustness, only studies containing at least 12 observations with complete information on disease incidence and yield were recorded. No minimum incidence range was imposed, and all recorded observations had disease incidence greater than zero. Yield values were standardized to a grain moisture content of 9% to ensure comparability among observations.

### Disease incidence–yield relationship

The relationship between peanut smut incidence and crop yield (kernel) was evaluated across studies to estimate disease-associated yield losses under field conditions. In all analyses, disease incidence was considered the explanatory variable and yield (kg ha⁻¹) the response variable.

The general linear damage function was defined such that the intercept (β₀) represents attainable yield under negligible disease pressure, whereas the slope (β₁) represents the damage coefficient, defined as the reduction in yield associated with each 1% increase in disease incidence.

### Exploratory segmented relationship

To explore potential deviations from linearity in the relationship between peanut smut incidence and yield, the pooled dataset was first examined using a locally weighted scatterplot smoothing (LOESS) function (Cleveland & Devlin, 1988), which suggested a possible change in slope at intermediate incidence levels (Fig. 3A). Based on the observed pattern, segmented regression models were fitted to test for potential breakpoints in the incidence–yield relationship (Muggeo, 2008). Model selection was based on visual inspection of the smoothed trend and comparative goodness-of-fit between linear and segmented formulations.

### Regression coefficients and meta-analytic modeling

#### Regression coefficients

Separate linear regression models were fitted to data from each study to describe the relationship between disease incidence (X, % infected pods) and yield (Y, kg ha⁻¹):

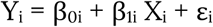

where β₀ᵢ represents the attainable yield in the absence of disease for the *i*-th study, β₁ᵢ represents the damage coefficient associated with each 1% increase in peanut smut incidence, and εᵢ represents the residual error term.

Study-specific regression coefficients were extracted to characterize variability in baseline productivity and disease-associated yield losses among environments (Madden et al., 2009ab; Paul et al., 2010; Barro et al., 2023).

#### Meta-analytic synthesis

Study-specific regression coefficients obtained from individual incidence–yield models were synthesized using random-effects meta-analytic models. Separate meta-analyses were conducted for intercepts (β₀) and slopes (β₁), allowing independent characterization of variability in attainable yield and disease-associated yield losses across environments (Duffeck et al. 2020; Madden and Paul 2009a).

For each study, the observed coefficient from each study (θi) was modeled as:

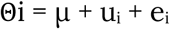

where μ represents the population-average effect, uᵢ the random deviation associated with the *i*- th study, and eᵢ the sampling error.

Random-effects models were fitted using restricted maximum likelihood (REML) as implemented in the ‘metafor’ package in R (Viechtbauer, 2010). Study-specific estimates were weighted by the inverse of their sampling variance, derived from the standard error of each regression coefficient.

Between-study heterogeneity was quantified using Cochran’s Q statistic, the inconsistency index (I²), and the between-study variance component (τ²), with τ representing the corresponding standard deviation (Higgins and Thompson, 2002).

The relative damage coefficient (DC) was calculated as:

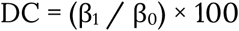

representing the percentage reduction in attainable yield associated with each 1% increase in peanut smut incidence (Lehner et al. 2016).

#### Moderator analysis

To explore potential sources of between-study heterogeneity, moderator analyses were conducted by stratifying studies according to disease incidence and yield levels (Madden and Paul, 2009b). Disease incidence categories were defined using the breakpoint estimated from segmented regression analysis of the incidence–yield relationship (Muggeo 2003, 2008). This threshold was used as an exploratory criterion and not interpreted as a biologically fixed cutoff. Low- and high-incidence categories were then incorporated as moderator variables to evaluate consistency in the incidence–yield relationship across strata.

#### Mixed-effects modeling

To account for the hierarchical structure of the dataset, in which observations were nested within studies, and to jointly estimate population-average and study-specific disease–yield relationships, linear mixed-effects models with random intercepts and random slopes were fitted (Barro et al. 2023; Gelman and Hill, 2007).

Yield observations (Yij) were modeled as a function of peanut smut incidence (Xij) according to:

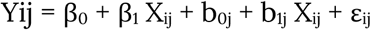

where β₀ and β₁ represent the fixed population-average intercept and slope, b₀ⱼ and b₁ⱼ represent study-specific random deviations in intercepts and slopes, and εᵢⱼ represents the residual error. Models were fitted by maximum likelihood using the lme4 package in R (Bates et al., 2015). This framework allowed simultaneous estimation of: (i) the average damage function across studies, (ii) variability in attainable yield among environments, and (iii) variability in disease-associated yield losses among studies.

## RESULTS

### Dataset overview

A high degree of variability was observed among studies in both peanut smut incidence and kernel yield. Peanut smut incidence ranged from 0.6% to 84.5%, while yield varied from 172 to 5,382 kg ha⁻¹ across observations. Substantial variability was observed across years. Incidence values spanned from less than 1% to more than 80% in multiple seasons, while yield distributions varied markedly among environments. General Deheza exhibited the widest range of both incidence (0.57 to 84.5%) and yield (172.4 to 4,932.2 kg ha⁻¹), as well as the largest number of observations (648 vs 274 from General Cabrera) (Fig. 1).

**Figure 1.**
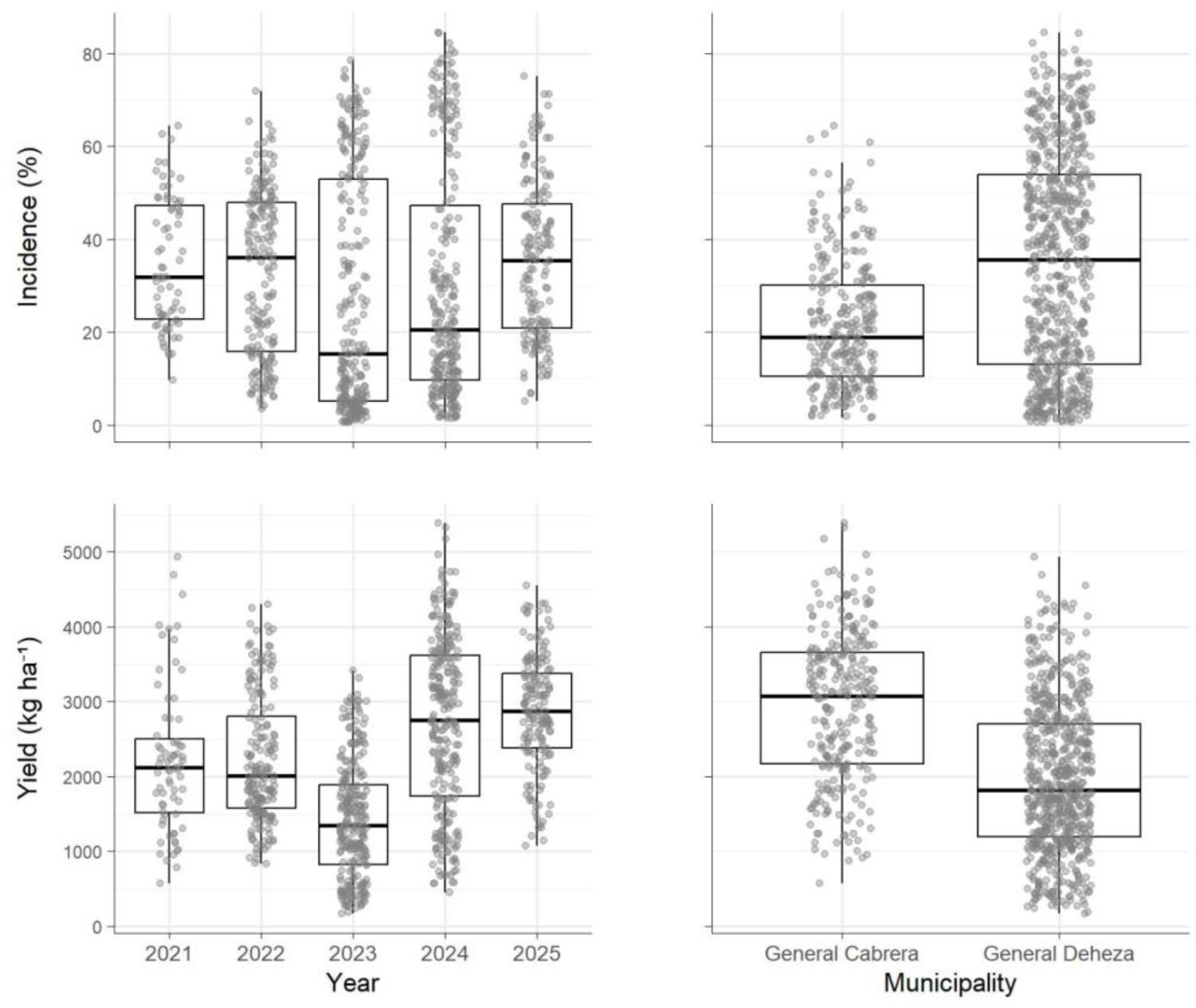
Distribution of peanut smut incidence and estimated kernel yield (kg ha⁻¹) across studies, years, and locations included in the meta-analysis. Grey dots represent individual observations corresponding to 1 m² sampling units.

### Global incidence-yield relationship

Across the complete dataset, peanut smut incidence was negatively associated with crop yield (Fig. 2). The global linear regression model estimated an average yield reduction of 26.9 kg ha⁻¹ for each 1% increase in disease incidence. The intercept, corresponding to the expected yield in the absence of disease, was estimated at 3,091 kg ha⁻¹. This relationship corresponds to an average reduction of approximately 269 kg ha⁻¹ for every 10 percentage-point increase in incidence. This estimate corresponds to a relative damage coefficient (DC) of 0.87%, indicating that each 1% increase in peanut smut incidence was associated with a yield reduction equivalent to 0.87% of the attainable yield under disease-free conditions.

**Figure 2.**
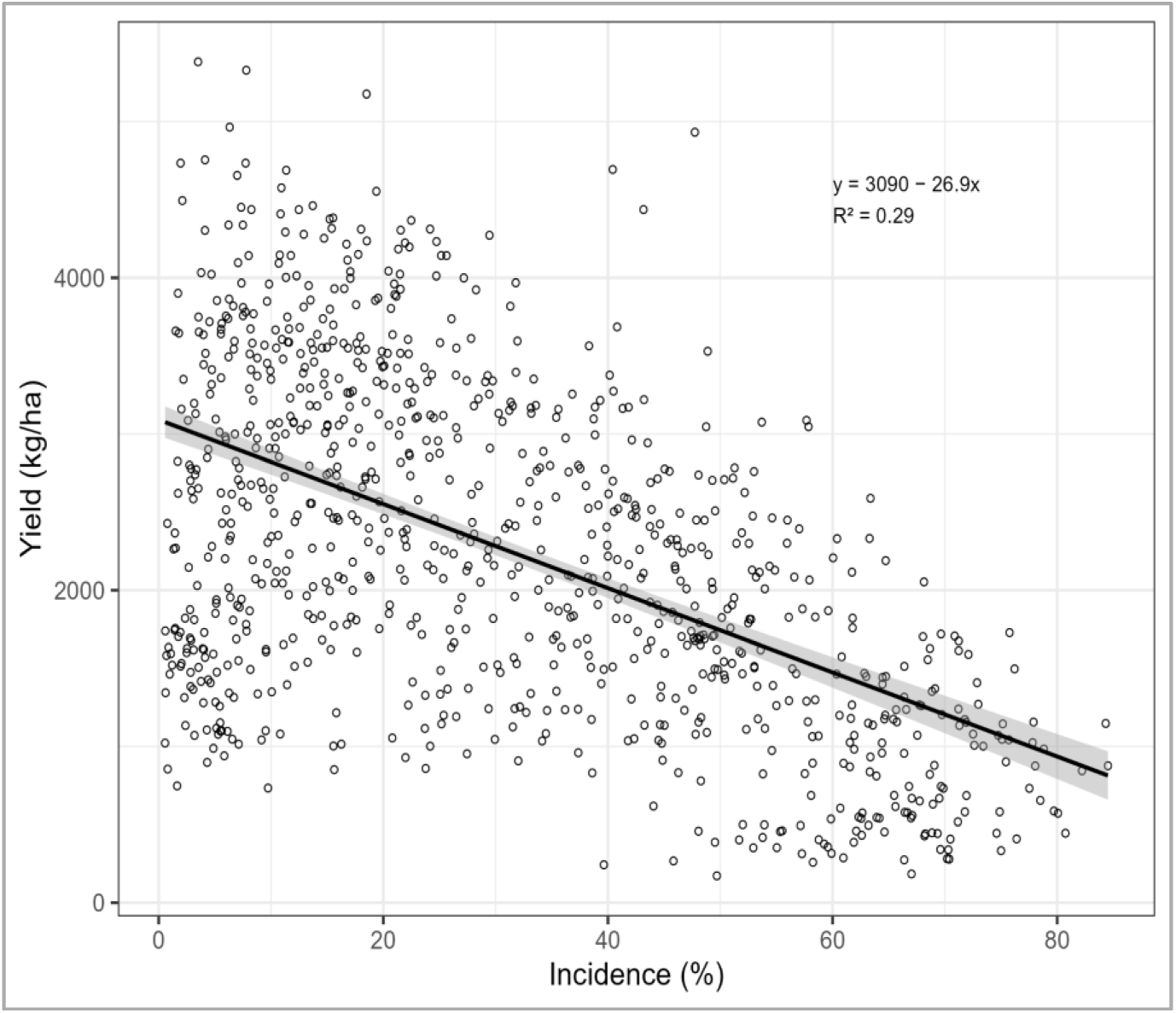
Global relationship between peanut smut incidence (%) and crop yield (kg ha⁻¹) across 922 observations from 26 field studies conducted in Argentina. Circlers represent individual sampling units. The solid line represents the fitted linear regression model, and the shaded area indicates the 95% confidence interval.

The model explained a low-to-moderate proportion of the total variability in yield (R² = 0.29), indicating that, although disease incidence was an important determinant of yield reduction, additional environmental and agronomic factors also contributed substantially to yield variability.

A greater dispersion of observations was observed at low incidence levels, where a wide range of yield values occurred under relatively limited disease pressure. Despite this variability, the overall negative trend remained consistent across the full incidence range, supporting a stable detrimental effect of peanut smut on crop productivity.

### Exploratory segmented relationship

The LOESS smoothing function suggested a possible change in slope at intermediate incidence levels (Fig. 3A). The segmented model identified a breakpoint at approximately 12.0% incidence (SE = 1.04), indicating a statistically significant change in slope across incidence levels (Fig. 3B). The segmented model provided a better statistical fit than the simple linear model (R² = 0.37 versus 0.29). Nevertheless, given the observational nature of the dataset and the strong heterogeneity among studies, the biological relevance of the identified breakpoint should be interpreted cautiously. Overall, these results suggest that the apparent non-linearity observed in pooled analyses is likely driven largely by between-study heterogeneity rather than by a true threshold response in the disease damage function.

**Figure 3.**
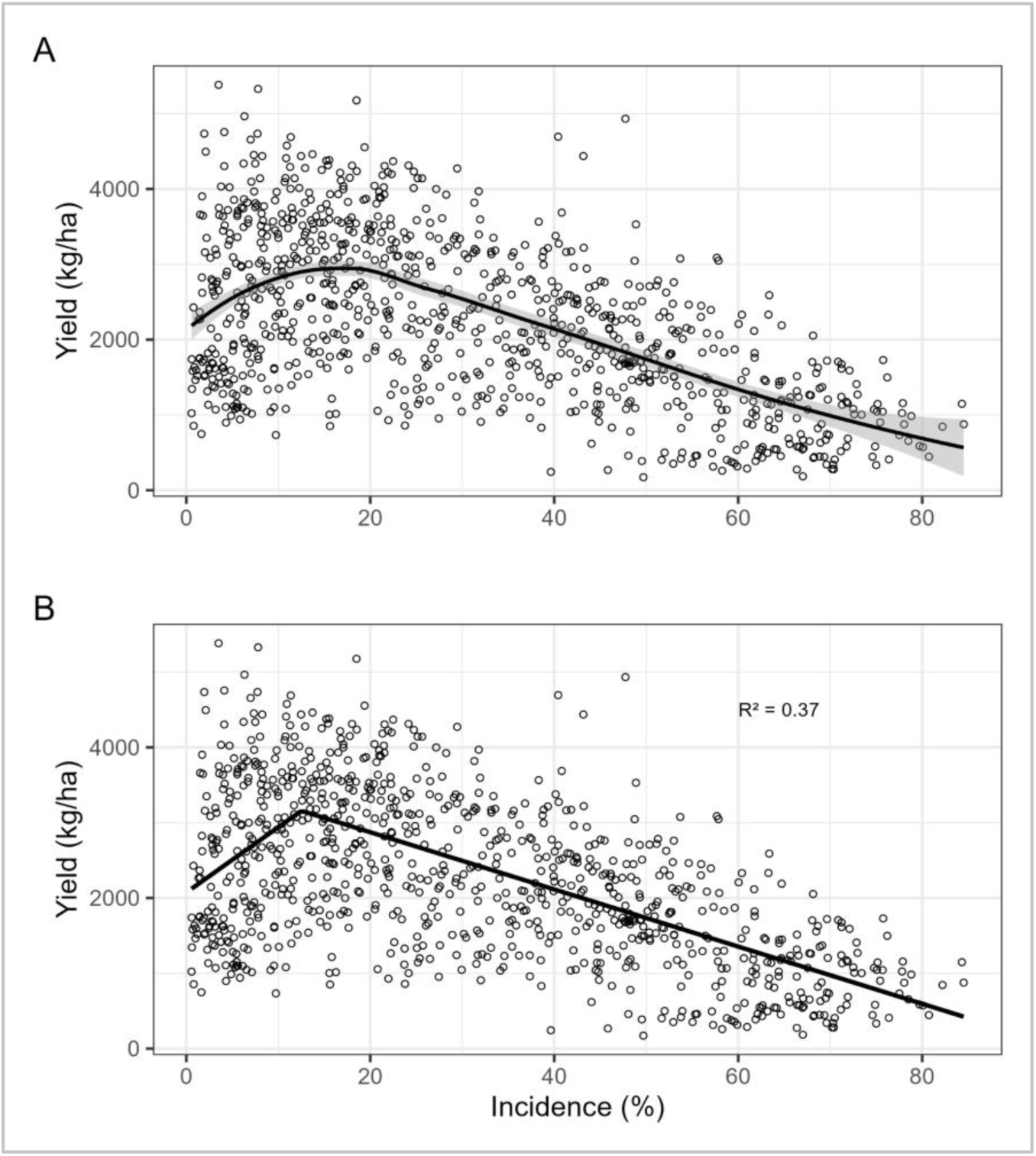
Exploratory assessment of non-linearity in the peanut smut incidence–yield relationship. (A) LOESS-smoothed relationship between peanut smut incidence (%) and crop yield (kg ha⁻¹). (B) Segmented regression fitted to pooled observations, identifying an exploratory breakpoint at approximately 12% incidence.

### Study-specific incidence–yield relationships

To further evaluate the consistency of the incidence–yield relationship across environments, linear regressions were fitted independently for each study. Considerable variability was observed among intercepts, reflecting substantial differences in attainable yield across experimental conditions. In contrast, slopes were comparatively consistent in both magnitude and direction. Most studies showed a negative association between peanut smut incidence and yield, whereas only a few exhibited weak or slightly positive slopes (Fig. 4A). These positive estimates were generally associated with studies containing predominantly low incidence values and are likely attributable to environmental or management-related factors rather than a biologically meaningful positive effect of the disease.

**Figure 4.**
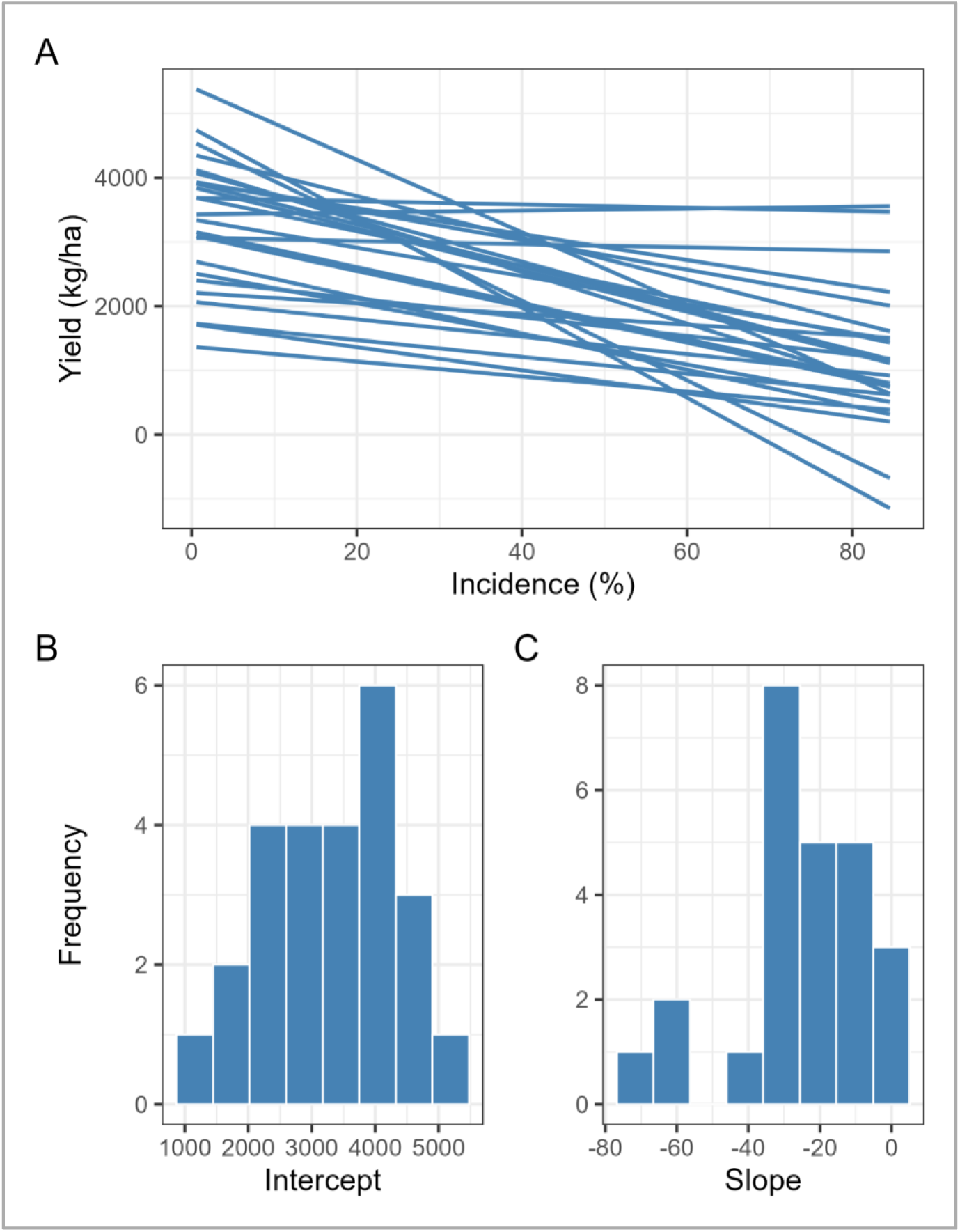
Study-specific incidence–yield relationships and variability in regression coefficients across studies. (A) Linear regression models fitted independently for each study (k = 26), describing the relationship between peanut smut incidence (%) and kernel yield (kg ha⁻¹). Each line represents a study-specific regression. (B) Frequency distribution of study-specific intercept estimates (β₀), representing attainable yield under negligible disease pressure. (C) Frequency distribution of study-specific slope estimates (β₁), representing the yield reduction (kg ha⁻¹) associated with each 1% increase in peanut smut incidence.

Expressed relative to attainable yield, study-specific damage coefficients were generally clustered around a common value, suggesting that disease effects were proportionally similar across environments despite large differences in baseline productivity. Visual inspection of the study-specific regression lines supported this interpretation, as most fitted lines differed primarily in vertical position rather than slope. Overall, these results indicate that much of the variability observed in pooled analyses was due to differences in attainable productivity across environments rather than to substantial changes in the underlying disease-damage relationship.

### Meta-analysis

Linear regressions between peanut smut incidence and yield were fitted independently for each study (k = 26), generating study-specific intercepts and slopes. Intercepts ranged from 1370 to 5409 kg ha⁻¹ (mean = 3331 kg ha⁻¹) (Fig. 4B), indicating substantial variability in attainable yield across environments. Slopes ranged from −70.1 to 6.7 kg ha⁻¹ per 1% increase in incidence (mean = −25.8) (Fig. 4C). Most studies exhibited negative incidence–yield relationships, although the magnitude of the estimated damage coefficient varied among studies (Fig. 5).

**Figure 5.**
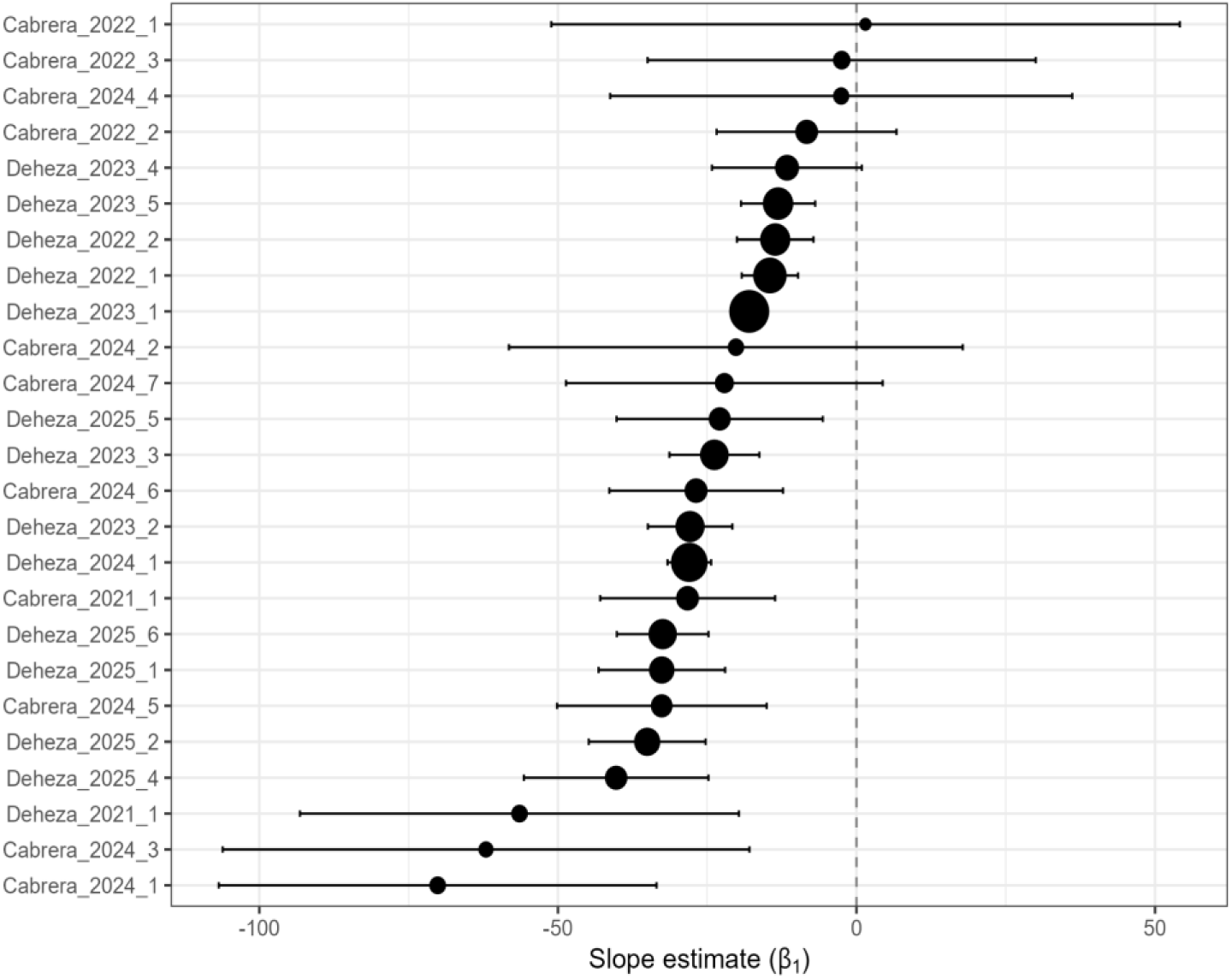
Forest plot showing study-specific slope estimates (β₁) obtained from linear regressions between peanut smut incidence (%) and crop yield (kg ha⁻¹) across independent studies (k = 26). Points represent study-specific estimates, and horizontal bars indicate 95% confidence intervals. Point size is proportional to study weight in the meta-analysis. Negative values indicate yield reduction associated with increasing peanut smut incidence. The vertical dashed line denotes the null effect (β₁ = 0), illustrating the predominance of negative disease-associated yield responses among studies.

The random-effects meta-analysis of intercepts estimated a population-average attainable yield of 3281 kg ha⁻¹ under negligible disease pressure (95% CI: 2985–3738; *p* < 0.001). However, heterogeneity among studies was extremely high (I² = 97.98%, Q = 1259.2, *p* < 0.001), indicating strong variability in baseline productivity across environments. The meta-analysis of slopes revealed a significant negative association between peanut smut incidence and yield, with an overall damage coefficient of −24.2 kg ha⁻¹ per 1% increase in incidence (95% CI: −29.0 to −20.6; *p* < 0.001). Given the estimated attainable yield of 3,331 kg ha⁻¹, this effect corresponds to a relative damage coefficient of approximately 0.74%, indicating that each 1% increase in disease incidence reduces yield by nearly 0.74% of the attainable yield. Although between-study heterogeneity remained substantial (I² = 75.2%), the direction of the effect was highly consistent among studies, with the vast majority of slope estimates remaining negative despite differences in magnitude (Fig. 5). To investigate whether the apparent non-linearity detected in segmented regression analyses reflected differences in damage coefficients across epidemiological contexts, incidence class was incorporated as a categorical moderator using the exploratory 12% threshold derived from the segmented model. This threshold was used exclusively as a data-driven stratification criterion. Incidence class did not significantly affect the slope estimates (QM = 0.01, *p* = 0.91) and explained none of the between-study heterogeneity (R² = 0%). Stratified models produced nearly identical incidence–yield relationships: studies below 12% incidence yielded the equation y = 3742.4 − 24.2x, whereas studies at or above 12% incidence yielded y = 3247.2 − 24.9x. Likewise, yield class (threshold = 2767 kg ha⁻¹) did not significantly affect slope estimates (*p* = 0.70). A marginal effect was detected for intercepts (*p* = 0.07), suggesting differences in attainable productivity among studies but not in the magnitude of disease-associated yield losses. Normalized yield response curves standardized to a common baseline further supported this interpretation, as curves corresponding to low- and high-incidence classes showed near-complete overlap (Fig. 6). Collectively, these results indicate that the apparent breakpoint observed in pooled analyses was primarily associated with variability in attainable yield among studies rather than with a true change in the underlying damage function.

**Figure 6.**
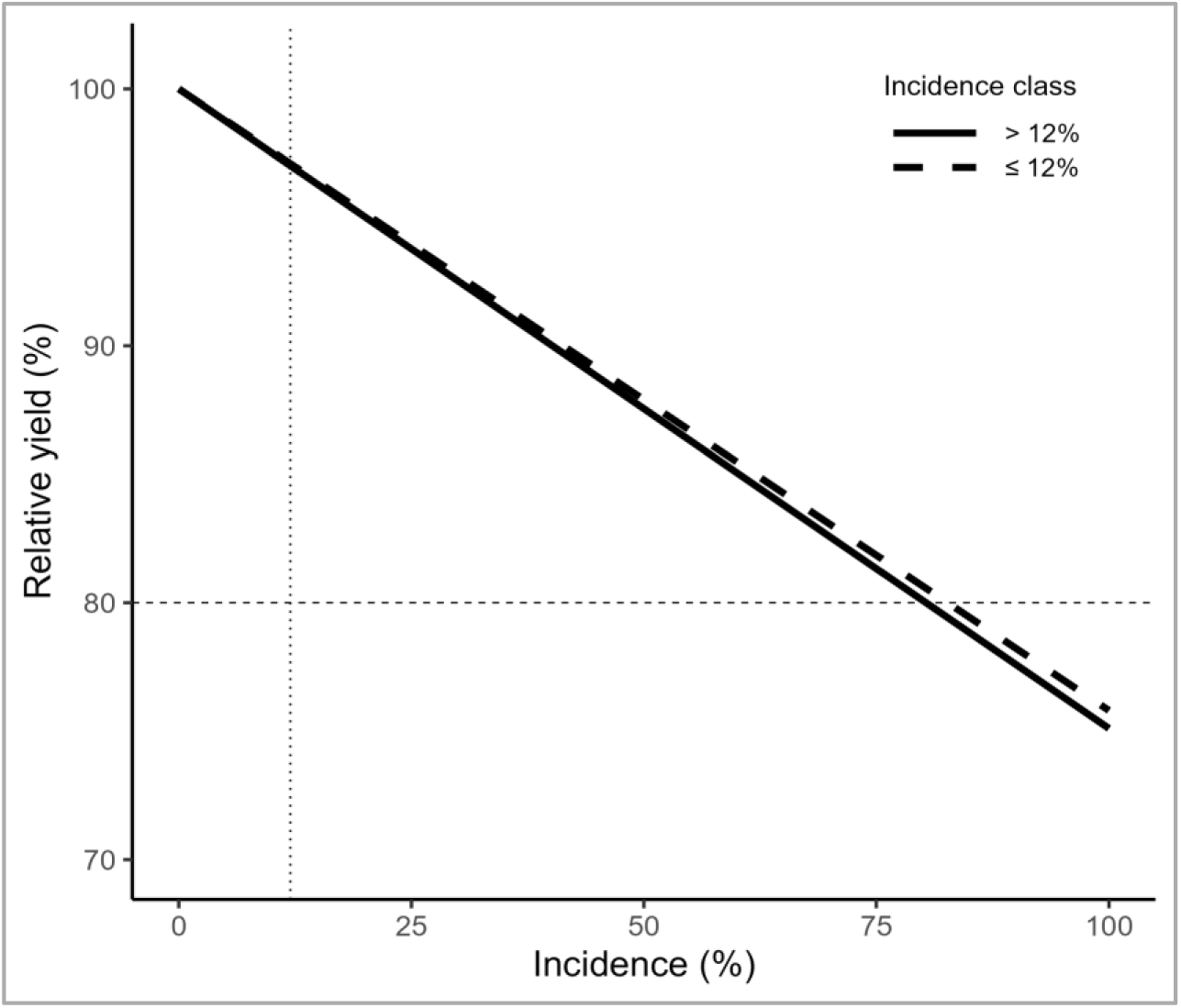
Standardized incidence–yield response curves stratified by incidence class. Normalized yield response curves standardized to a common baseline (100% relative yield at zero disease incidence) for studies classified according to the exploratory incidence threshold derived from segmented regression (≤12% and >12% smut incidence).

### Mixed-effects modeling

Linear mixed-effects models with random intercepts and random slopes confirmed the overall negative relationship between peanut smut incidence and yield. The population-average model estimated an attainable yield of 3,375 kg ha⁻¹ under negligible disease incidence and an average kernel yield reduction of 28.7 kg ha⁻¹ for each 1% increase in incidence. This relationship corresponds to a relative damage coefficient of approximately 0.85%, which was remarkably similar to estimates obtained using the other analytical approaches. Substantial variability among studies was observed for intercepts (SD = 874 kg ha⁻¹), indicating marked differences in attainable productivity across environments. In contrast, variability in slopes was comparatively small (SD = 6.2 kg ha⁻¹ per 1% incidence), suggesting that disease-associated yield losses were relatively stable among studies. Overall, the mixed-effects framework reinforced the conclusions obtained from the meta-analytic approach, indicating that most between-study heterogeneity was associated with differences in baseline productivity rather than major shifts in the incidence–yield damage relationship.

Across analytical approaches, attainable yield estimates ranged from 3,091 to 3,375 kg ha⁻¹, whereas damage coefficients varied from 24.2 to 28.7 kg ha⁻¹ per 1% increase in incidence (Table 1). When expressed relative to attainable yield, estimated damage coefficients ranged from 0.74 to 0.87%. Despite substantial heterogeneity in attainable productivity among studies, the proportional effect of peanut smut incidence on yield remained remarkably stable across models.

**Table 1.**
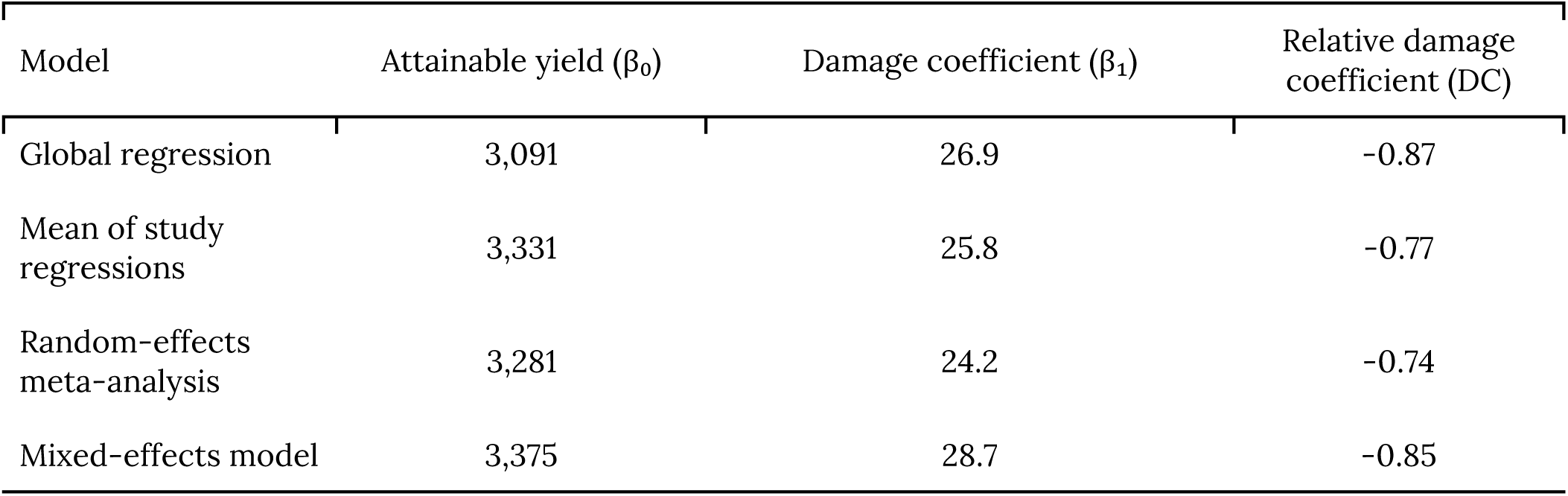
Summary of regression-based estimates of attainable peanut kernel yield (β₀; kg ha⁻¹), damage coefficient (β₁; kg ha⁻¹ yield loss per 1% increase in peanut smut incidence), and relative yield loss (% per 1% incidence) obtained from different analytical approaches.

## DISCUSION

The present study provides the first quantitative synthesis of the relationship between peanut smut incidence and yield loss across field environments in Argentina, demonstrating a consistent and significant negative association between disease intensity and crop productivity. Across analytical approaches, peanut smut incidence was associated with substantial reductions in yield, with population-average estimates ranging from approximately 25 to 29 kg ha⁻¹ of peanut yield kernels for every 1% increase in disease incidence. When expressed relative to attainable yield, these estimates corresponded to damage coefficients ranging from 0.74 to 0.87%, indicating that each 1% increase in disease incidence reduced yield by approximately three-quarters of one percent of the attainable yield. These findings quantitatively support previous observational evidence indicating that peanut smut still represents one of the most economically damaging diseases affecting peanut production in Argentina (Paredes et al., 2024; Rago et al., 2017) and reinforce the importance of effective disease management strategies under field conditions (Cazón and Paredes, 2026).

A central finding of this study was the relative stability of the disease-associated damage coefficient across environments despite substantial variation in attainable yield. Regardless of the analytical approach employed, relative damage coefficients remained remarkably consistent, ranging from 0.74 to 0.87%. This narrow range contrasts sharply with the large variability observed in attainable kernel yield estimates and indicates that peanut smut exerts a proportionally similar effect on crop productivity across a wide spectrum of production environments. Visual inspection of the pooled dataset initially suggested a nonlinear relationship between peanut smut incidence and yield, characterized by an apparent change in slope around 12% incidence. Because tolerance and compensatory responses at low disease levels have been reported in some crop–pathogen systems, where physiological and developmental adjustments can partially mitigate the impact of disease on crop performance (Bingham et al., 2009; Willocquet & Savary, 2026), this apparent threshold warrants further investigation. Although segmented regression yielded a statistically better fit than a simple linear model, subsequent analyses did not support the presence of a biologically meaningful breakpoint. Study-specific regressions, moderator analyses, and mixed-effects modeling consistently indicated that the incidence–yield relationship remained remarkably stable across studies, with no significant differences in disease-associated damage coefficients among incidence classes. Likewise, normalized response curves exhibited substantial overlap throughout the incidence range.

In contrast, substantial variation was observed in model intercepts, reflecting marked differences in attainable productivity among environments. This pattern suggests substantial heterogeneity in attainable yield among studies rather than major changes in the functional form of the disease– yield relationship. Accordingly, the apparent nonlinearity observed in the pooled dataset does not reflect changes in the biological effect of peanut smut on yield, but rather differences in baseline productivity among studies. Because environments with contrasting yield potential contribute observations across different incidence ranges, pooling data without adequately accounting for study-level variability can generate apparent threshold responses that are statistical artifacts rather than genuine epidemiological phenomena (Madden and Paul, 2009a; Barro et al., 2023). This interpretation is further supported by conceptual and empirical studies showing that variation in attainable yield can strongly influence the shape of observed damage functions without necessarily altering the underlying disease-associated damage coefficient (Willocquet and Savary, 2026).

Consistent with this interpretation, mixed-effects analyses revealed considerable heterogeneity in intercepts but comparatively limited variation in slopes. Therefore, peanut smut appears to reduce yield at a relatively constant rate across production environments, while environmental and agronomic conditions primarily determine the productivity level from which these losses occur (Madden et al., 2007; Willocquet and Savary, 2026). This pattern is also consistent with the highly aggressive nature of peanut smut in susceptible cultivars, where infection typically results in a high proportion of severely damaged pods, with limited variability in symptom expression among susceptible genotypes and across disease pressures (Paredes et al., 2022, 2024).

The strong heterogeneity observed for attainable yield was expected given the diversity of environmental and agronomic conditions represented in the database. Yield formation in peanuts is strongly influenced by climatic conditions and seasonal variation, which can vary substantially among locations and years (Duncan et al., 1978; Wei et al., 2022). Consequently, environments with contrasting productive potential may exhibit similar disease effects in relative terms while differing markedly in absolute yield. The very high heterogeneity observed for intercepts, together with comparatively lower variability in slopes, supports the interpretation that peanut smut exerts a relatively stable penalty on productivity irrespective of environmental context, whereas environmental and agronomic factors primarily determine the attainable yield from which these losses occur (Barro et al., 2023; Willocquet and Savary, 2026).

From an applied perspective, the identification of a relatively stable damage coefficient has important implications for disease management and loss estimation. The estimated yield penalty of approximately 240–290 kg ha⁻¹ for every 10 percentage-point increase in incidence provides a biologically interpretable and operationally useful metric for estimating potential economic losses under field conditions. Expressed on a relative basis, the estimated damage coefficients indicate kernel yield reductions of approximately 7–9% for every 10 percentage-point increase in incidence, depending on the analytical framework considered. Such estimates may support disease monitoring programs, assist in evaluating the economic benefit of management practices, and contribute to decision-making regarding cultivar selection, crop rotation, and integrated disease management strategies (Paredes et al., 2024). Furthermore, the relative consistency of the disease–yield relationship suggests that incidence measurements, despite their simplicity, may provide sufficient information for predicting yield penalties across a broad range of epidemiological scenarios. The use of incidence as the primary disease descriptor is also biologically justified in the peanut smut pathosystem. Although disease severity can be quantified through pod damage scales, Paredes et al. (2022) reported that approximately 80% of infected pods correspond to the most severe damage categories. This predominance of severely affected pods reduces the additional discriminatory value of severity assessments and supports the use of incidence as a robust and operationally efficient indicator of disease impact. Under these conditions, the proportion of infected pods may better represent the overall burden of disease at the field level while remaining practical for large-scale monitoring and yield-loss estimation.

The methodological framework adopted in this study also highlights the value of meta-analytic and mixed-effects approaches for understanding disease–yield relationships in pathosystems characterized by high environmental variability. Previous syntheses in wheat, soybean, potato, and other crop systems have demonstrated the importance of separating true disease effects from environmental noise when estimating yield losses (Barro et al., 2021; Bingham et al., 2009; Duffeck et al., 2020; Yellareddygari et al., 2018). By explicitly modeling between-study heterogeneity, the present study extends this framework to peanut smut and provides a population-level estimate of disease-associated losses while preserving information on environmental variability.

Some limitations should nevertheless be acknowledged. The database was assembled from observational field studies originally designed for disease management evaluation rather than for formal yield-loss experimentation, and therefore residual confounding associated with environmental or management variables cannot be entirely excluded. In addition, the dataset was geographically concentrated in the core peanut-producing region of Argentina, which may limit extrapolation to emerging production areas with distinct inoculum histories or environmental conditions. Future studies incorporating additional regions, cultivars, and epidemiological descriptors, including disease severity, severely damaged pods, or temporal disease development, may further refine predictive models of peanut smut-associated yield loss.

Overall, our findings indicate that peanut smut consistently reduces peanut yield across production environments, with relative damage coefficients converging around 0.8% yield loss per 1% increase in disease incidence. This quantitative characterization of the incidence–yield relationship provides an epidemiological basis for estimating economic losses and contributes to the development of evidence-based management strategies for one of the most important diseases affecting peanut production in Argentina.

## Acknowledgements

We wish to thank INTA and Fundación Maní Argentino for providing resources for compiling this project.

## Author’s contribution

LIC and JAP conceptualized the study, wrote the manuscript, and conducted the data analysis. EMDP and ACCC contributed to the data analysis, revision, and co-wrote the manuscript. NRG, FA, and BXC contributed to data compilation and manuscript revision.

## Data Availability

The datasets generated and/or analyzed during the current study are available at the following repository link: https://github.com/ignaciocazon/Peanut-smut-Meta-anlysis

## Funding

This work was supported by INTA [Project I090, 2023-PD-L01-I074] and Fundación Maní Argentino

## Declarations Conflict of interest

All authors declare that they have no conflicts of interest.

## Notes

https://github.com/ignaciocazon/Peanut-smut-Meta-anlysis

### Competing Interest Statement

The authors have declared no competing interest.

